# Nucleosome positioning on large tandem DNA repeats of the ‘601’ sequence engineered in *Saccharomyces cerevisiae*

**DOI:** 10.1101/2021.08.25.457076

**Authors:** Astrid Lancrey, Alexandra Joubert, Evelyne Duvernois-Berthet, Etienne Routhier, Saurabh Raj, Agnès Thierry, Marta Sigarteu, Loic Ponger, Vincent Croquette, Julien Mozziconacci, Jean-Baptiste Boulé

## Abstract

The so-called 601 DNA sequence is often used to constrain the position of nucleosomes on a DNA molecule in vitro. Although the ability of the 147 base pair sequence to precisely position a nucleosome in vitro is well documented, in vivo application of this property has been explored only in a few studies and yielded contradictory conclusions. Our goal in the present study was to test the ability of the 601 sequence to dictate nucleosome positioning in *Saccharomyces cerevisiae* in the context of a long tandem repeat array inserted in a yeast chromosome. We engineered such arrays with three different repeat size, namely 167, 197 and 237 base pairs. Although our arrays are able to position nucleosomes in vitro as expected, analysis of nucleosome occupancy on these arrays in vivo revealed that nucleosomes are not preferentially positioned as expected on the 601-core sequence along the repeats and that the measured nucleosome repeat length does not correspond to the one expected by design. Altogether our results demonstrate that the rules defining nucleosome positions on this DNA sequence in vitro are not valid in vivo, at least in this chromosomal context, questioning the relevance of using the 601 sequence in vivo to achieve precise nucleosome positioning on designer synthetic DNA sequences.

## Introduction

The nucleosomes are more than structural proteins that enable folding of a meter-long molecule into a micrometer-diameter nucleus, and are now recognized as master players in the many facets of genome metabolism. The deposition of post-translational modifications on nucleosomes along genomes are tightly correlated with the regulation of gene expression [1]. The first experimental evidence for the existence of nucleosomes came from their regular, periodic assembly along the DNA molecule [2, 3]. In the following decades, this periodicity was measured in various contexts and the Nucleosomal Repeat Length (NRL) was shown to vary in different species, different cell types and even in different regions of chromosomes [4]. As more measures became available, Jonathan Widom noticed that the NRL took preferential values of 167 + *n* ∗ 10 base pairs (bp) and linked this observation to the value of the DNA helical pitch [5]. As the periodic position of nucleosomes became of high mechanistic interest to explain chromatin function, nucleosomal array reconstruction protocols for in vitro studies were developed. In order to test the physical and structural properties of DNA wrapping around an histone octamere and the role of the underlying DNA sequence, Widom and colleagues isolated, using a SELEX approach, a set of DNA sequences that could precisely position nucleosomes in vitro [6]. Among these 147 bp DNA sequences, the ”601” sequence was retained to be the one forming the most stable nucleosomal complex in vitro, together with an histone octamere. This ”601” sequence has been successfully used in many in vitro studies to reconstruct regular arrays of nucleosomes for electron microscopy [7, 8] and single molecule manipulations [9–11]. These studies confirmed that the 167 + *n* ∗ 10 bp NRL rule was required to form regular arrays and revealed both their ambivalent ability to be highly resilient to supercoiling in their extended form [9, 12] and to fold into very dense and compact fibers under specific buffer conditions [13]. The detailed biophysical modeling of nucleosomal array structures showed that the observed quantized behavior of the NRL is primarily due to the combination of the nucleosome/nucleosome steric hindrance and the 10 bp helical pitch of the DNA linker [14, 15].

The potential use of the 601 sequence for in vivo studies has early been envisioned by Lowary and Widom themselves. However, only a handful of studies have been carried so far in this direction and the ability of one inserted sequence to position a nucleosome has led to divergent conclusions that pointed to the dependence on the context in which the 601 sequence was used (episomal vs genomically inserted) [16–19]. The possibility of forming a regular array of nucleosomes using these positioning sequences in cells has been attempted in yeast but the positioning of nucleosomes on these arrays has not been firmly established [20]. Our goal here was to assess the ability of 601 sequence to position nucleosomes in vivo on long tandem arrays using three different linker lengths of 20, 50 and 90 bp, that would respectively constrain the NRL respectively to 167, 197 or 237 bp. Using a genome editing approach that we recently developed to engineer long DNA tandem repeats in yeast chromosomal DNA [21], we assembled tens of 601 repeats and analyzed the statistical positioning of nucleosomes on these arrays. Our results show that, in vivo, the 601 sequence is not able to position the nucleosomes on the 601 core, yielding NRL which are different from the ones we designed, except for the control 167 bp tandem array which correspond to the naturally observed yeast NRL.

## Results

### In vivo assembly of 601 tandem DNA repeats in the yeast genome

We used a strategy that we developed recently [21] to assemble tandem DNA repeats containing the 601 sequence in the genome of *Saccharomyces cerevisiae*. Specifically, we engineered three designs of overlapping oligonucleotides to partially replace the non-essential YMR262 gene and its promoter, located in chromosome XIII, to give rise to arrays of tandem repeats of 167, 197 or 237 bp long monomers (Figure 1 and supplementary Figure S1, S2). Integration at the YMR262 locus resulted in the removal of 129 base pairs upstream the start codon and 231 bp downstream the start codon [21]. Sizes of arrays obtained by this approach ranged from one repeat to arrays longer than 15 *kb*, as estimated by gel electrophoresis, corresponding to more than 50 repeats (Figure 2). Correct assembly was verified by sequencing of the junctions (Supplementary Figure S2). For each of the three designs, the clone with the largest array was kept for chromatin studies. Since tandem repeats may be susceptible to genomic instability, we measured the stability of the arrays upon mitotic division. After around 25 generations, only few rearrangements were observed. The percentage of rearranged clones was respectively 1, 2 and 4%, demonstrating that large tandem repeats of the 601 sequences are easily maintained in the yeast genome (Supplementary Figure S3).

**Figure 1.**
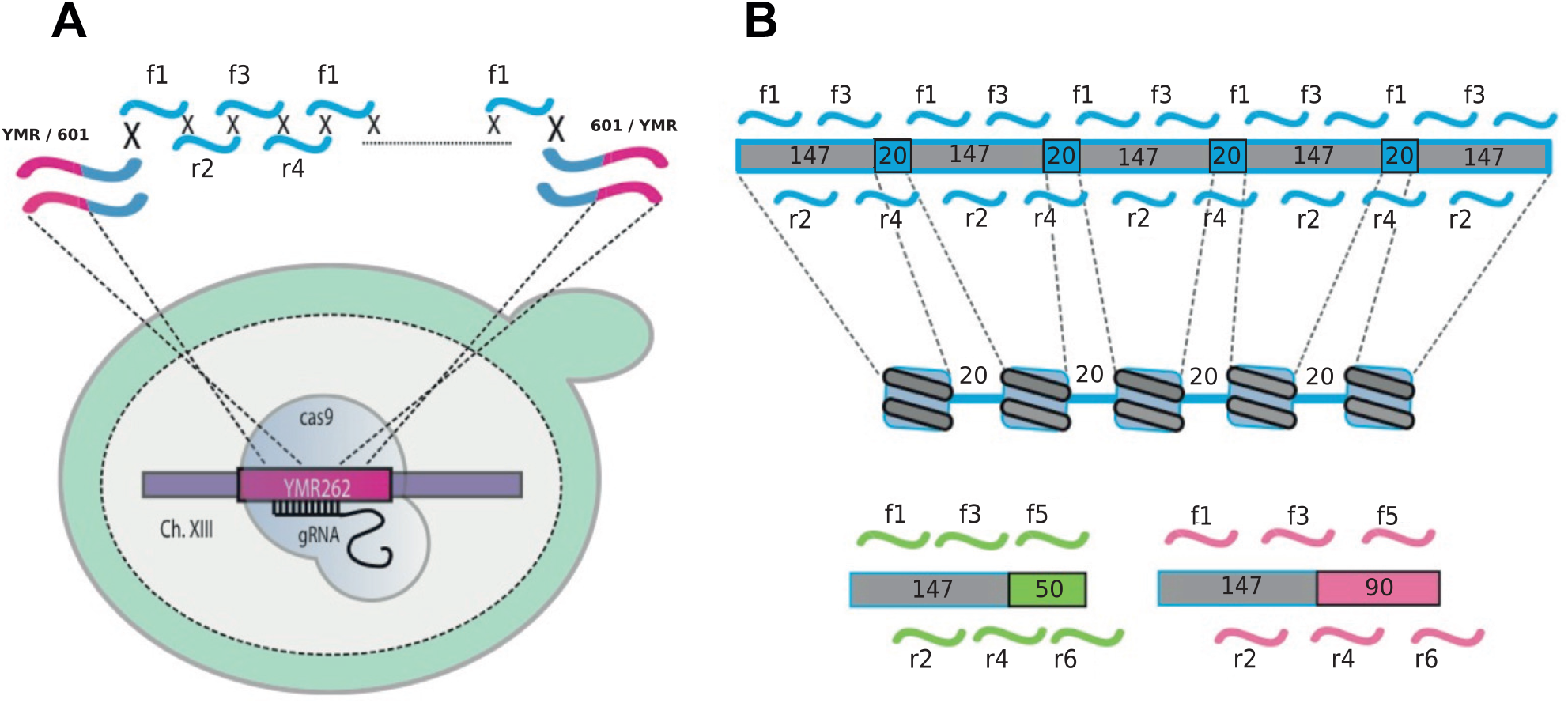
In vivo assembly of ‘601’ tandem DNA repeats. (A) Schematic representation of the CRISPR/Cas9-assisted genomic integration of overlapping oligonucleotides. The method is represented for integration of 601-167 repeats in the YMR262 gene of *S. cerevisiae*. (B) Design of the 601-167 repeats: four 50 nt oligonucleotides f1-r2-f3-r4 overlapping on 20 nt were used to construct the 601-167 array. The 147 bp core sequence (grey) corresponds to the DNA portion theoretically wrapped around one nucleosome, followed by a 20 bp linker (blue). The f4 oligo overlaps with f1 so that several monomers can assemble into a repeated array. (C) Design of the 601-197 and 601-237 arrays, using 6 six overlapping oligonucleotides.

**Figure 2.**
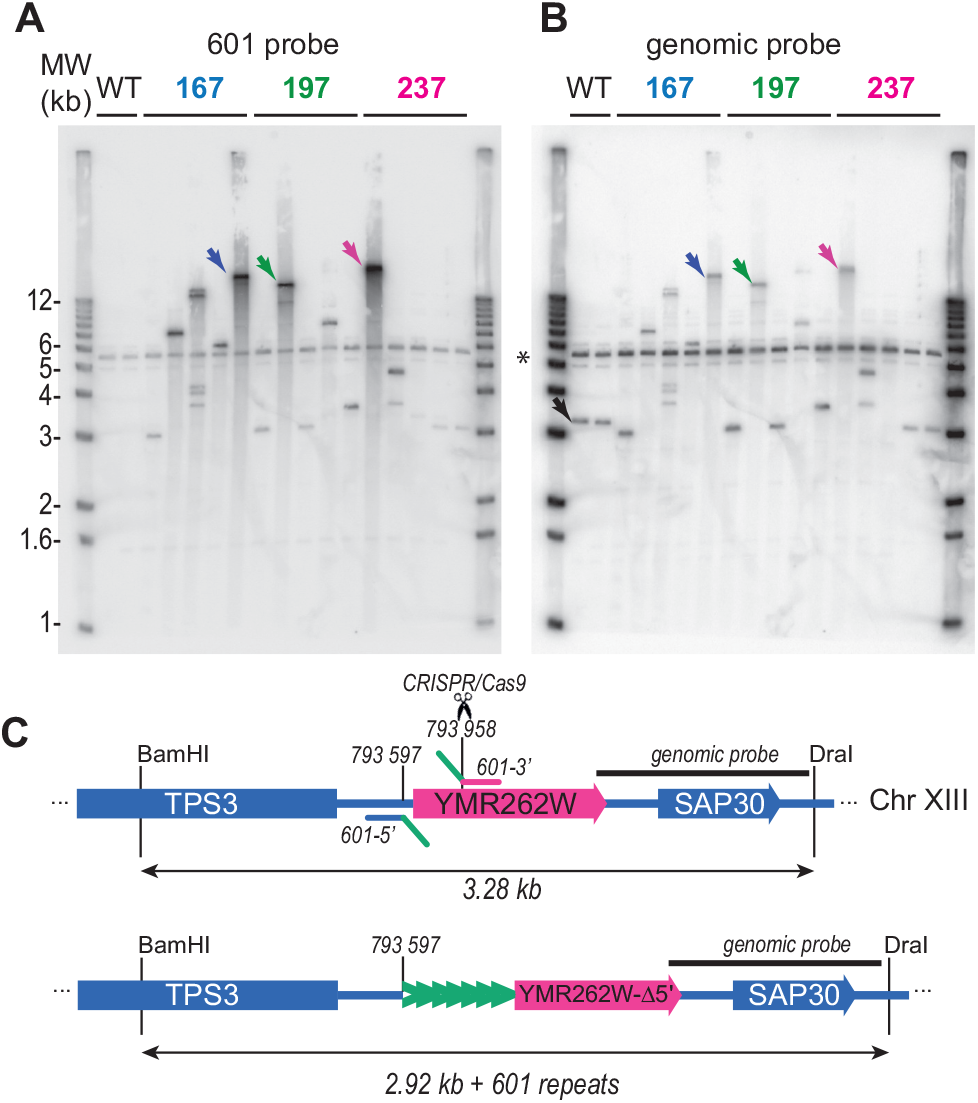
Overlapping oligonucleotides assemble to form long repeated arrays at the YMR262 locus. (A) Diagram of the assembly region before and after CAS9-assisted recombinational assembly. Position of the genomic probe is represented by a dark line. Localization of the genomic-601 junction primers are also indicated. (B) Southern Blot analysis of yeast recombinant strains. Genomic DNA from strains containing 601 repeats were digested with BamhI and DraI. Samples were electrophoresed, blotted and hybridized with a 601 probe containing a 601-147 nt repeat unit. The first two lanes (WT) correspond respectively to the wild-type and CAS9 expressing strains. Five clones are shown for respectively the 167, the 197 and the 237 design; the strains containing arrays largers than 15 kb (marked with color arrows) were retained for further manipulations. (C) The membrane shown in B was stripped and rehybridized using the genomic probe shown in A. The star marks a non specific band present on both Southern blots and present in the WT controls

### The NRL measured on 601 repeats are different in vitro and in vivo

In order to verify that our design of the DNA linkers yielding the 167, 197 and 237 bp repeats did not affect the ability of the 601 sequence to position precisely nucleosomes in vitro, we assembled chromatin on plasmids containing respectively 20, 14 and 14 repeats of the 167, 197 and 237 bp repeats (supplementary Figure S4A-C). In vitro assembled chromatin was digested by MNase (Micrococcal Nuclease), separated on gel and probed by Southern Blotting using a 601 probe. The results confirmed that nucleosomes were spaced as expected on all three 601 repeats (supplementary Figure S4D, E). However with the 601-237 design we observed two additional shorter ladders reflecting that some nucleosomes are positioned according to a second NRL when the designed NRL is equal to 237 bp.

To get insights into nucleosome organization in the 601 repeats *in vivo* we first performed MNase-gel experiments on the wild type YPH499 strain and the three yeast strains harboring the repeats. Mnase-gel experiments consisted on partially digesting the chromatin of cells harboring 601 repeats with Mnase, followed by gel analysis of NRL over the whole genome or specifically within the repeats (Figure 3, see Material and Methods for details). We measured a NRL of 163 ±1 bp for the whole genome, similar to the values expected for the mean NRL in *S. cerevisiae* genome [22]. The values measured after Southern blotting within the 601 arrays revealed that nucleosomes are spaced in average according to a 170±4 bp NRL, a 176±7 bp NRL and a 171*±* 9 bp NRL respectively in the 167 bp, the 197 bp and the 237 bp repeats (Figure 3C,D). The NRL measured for the 601-167 repeats is close to the one expected, however nucleosomes did not appear preferentially spaced according to the pre-defined 197 and 237 bp NRL in the respective 601-197 and in the 601-237 design.

**Figure 3.**
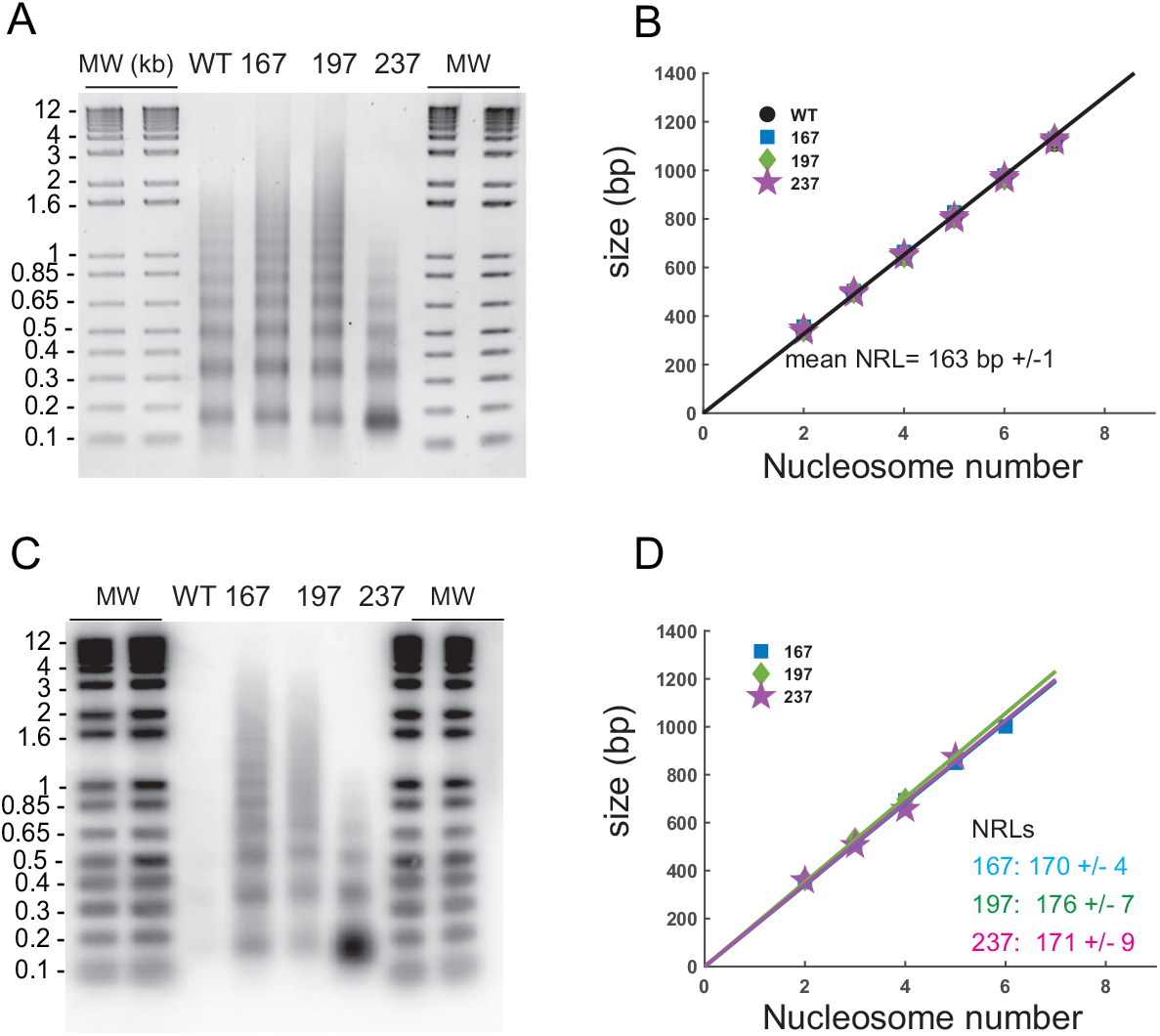
NRL estimation in ‘601’ tandem DNA repeats in vivo. (A) MNase gel on which were migrated the purified DNA from partial MNase digestions of the wt and the three 601 strains chromatin. Each band from one nucleosomal ladder corresponds respectively, from bottom to top, to the mononucleosomal DNA, di-tri-, etc.. Conversion of migration distance inside the gel into DNA size enabled to determine the DNA size of each nucleosomal band of the ladders and to calculate the expected whole genome average 163 bp NRL of the WT strain (B). (C) The EtBr gel in (A) was transferred onto a membrane and hybridized with the 601 radio-labeled probe to specifically determine the NRL in the 601 repeats. (D, E, F) Results of NRL calculation in the 601 repeats for respectively the 167, 197 and 237 strain.

### Nucleosomes are not positioned on the 601 core sequence in vivo

To have a more precise assessment of nucleosome occupancy in the repeats *in vivo*, we sequenced mononucleosomal DNA from the 601 strains and realigned all assigned midpoint of MNase resistant fragments on a single 601 repeat sequence (see material and methods for details and Supplementary Figure S5, S6). We used two levels of MNase digestion: moderate and over-digested, with two replicates for each conditions. Nucleosomal DNA midpoint density on the three array types are presented on Figure 4A-C. The length distribution of moderately digested fragments was found to be centered on 147 bp (coral and purple distributions on Figure 4D-F) whereas the length distribution of over-digested fragments was closer to 125 bp, exhibiting two to three peaks corresponding to various levels of digestion of the DNA wrapped around the nucleosome (green and cyan distributions on Figure 4D-F). For over-digested chromatin, we observed a main peak in the distribution of fragments midpoints, at position 110 on the 167 bp repeats, and at position 150 and 160 on respectively the 197 and 237 bp repeats (Figure 4 green and cyan). Regardless of the level of Mnase digestion, density patterns of fragments midpoints gave similar peak positions, although patterns obtained with over digested fragments gave a stronger main peak for the 167 and 197 repeats (Figure 4 coral and purple curves).

**Figure 4.**
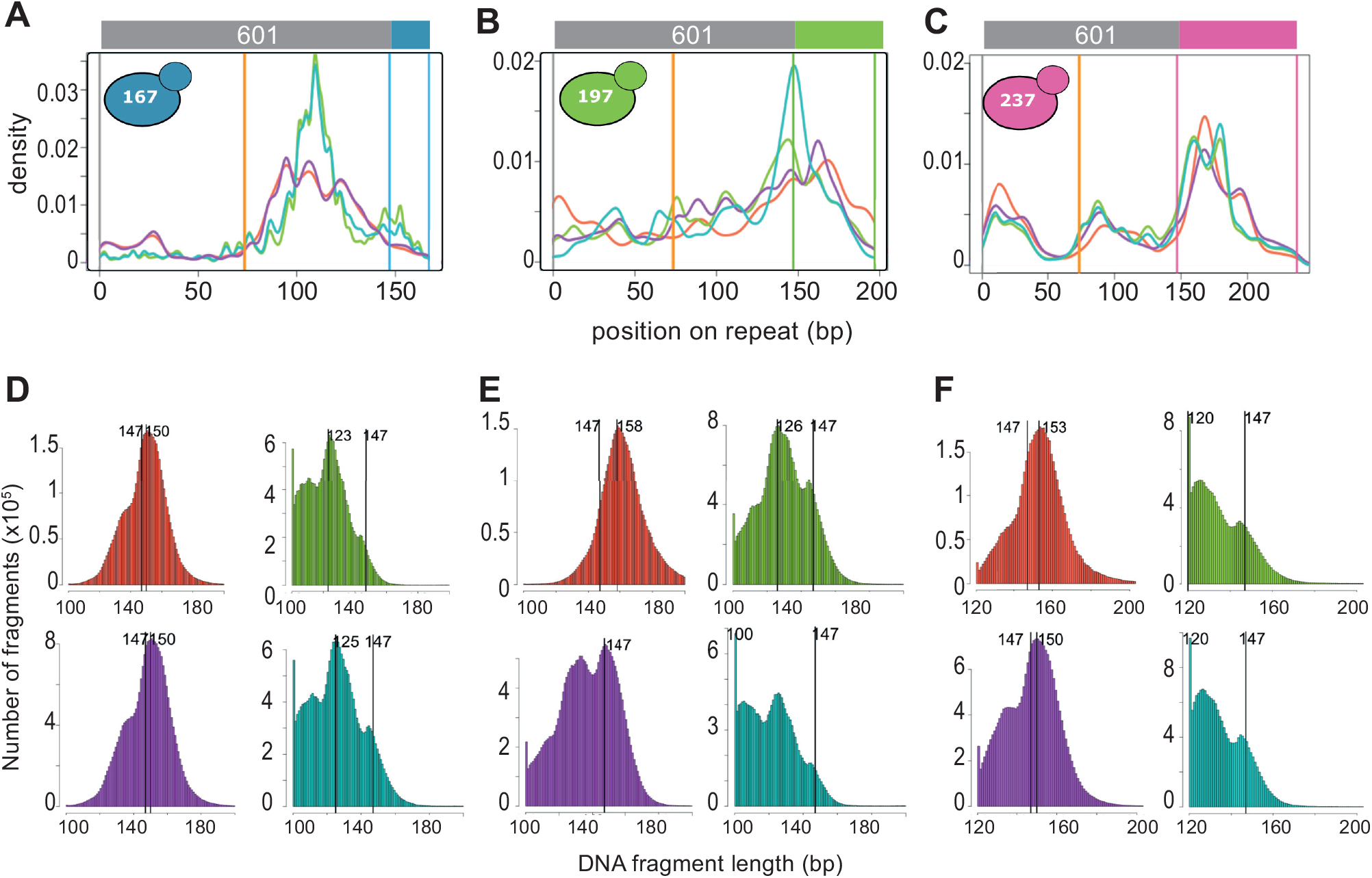
Nucleosomal fragment midpoints are not localized at the middle of 601 cores in the repeats *in vivo*. Mid-points Kernel density curves of each individual library are displayed on the same graphic for each 601-strain: the 601-167 strain (A), the 601-197 strain (B) and the 601-237 strain (C). The orange vertical line is localized at the 601-core middle and the two other colored vertical lines delimit the linker portion of the corresponding 601 design. The fragment length distribution histograms of the four corresponding libraries are displayed for the 601-167 strain (D), the 601-197 strain (E) and the 601-237 strain (F).

Altogether, our results indicate that nucleosomes assembled on the DNA repeats are not wrapped around the 601 core DNA according to in vitro derived rules, since none of the detected peaks are centered on the 601 core. Surprisingly, we found that nucleosomes tend to position themselves on the linker part of the design rather than on the 601 core.

For each the three arrays tested, the positioning of the nucleosome center on the 601 repeat is very reproducible and does not depend on the digestion level. For the 167 bp repeats most of the nucleosomes are positioned around the middle of the second half of the 601 core sequence (Figure 4A). For the 197 bp repeats, the positioning is more uniform, with a higher density at the boundary between the 601-core end and the linker (Figure 4B). For the 237 bp repeats, we detected three preferential positions centered respectively around 20, 90 and 160 bp(Figure 4C). In summary, nucleosomes are periodically positioned according to the pre-defined NRL only for the 601-167 repeats. However they are not preferentially found on the 601 core sequence as expected. In the two other 601 designs we did not observe an exclusive nucleosome spacing according to the designed NRL, a results which confirms our MNase-gel results (Figure 3E and F). For these two designs, we find as well that the 601 core sequence is more repleting than attracting for nucleosomes.

### In vivo assembled 601 repeats are weakly transcribed

Since the transcription machinery is strongly associated with chromatin remodeling complexes that could have a predominant role in nucleosome occupancy, we then checked the transcription activity of our synthetic 601 arrays. To see whether deleting the YMR262 promoter upon repeat assembly was sufficient to abolish polymerase II transcription through the assembled array and the remaining of the YMR262 gene, we measured genome wide transcription in exponentially growing cells by RNA-seq. We then assessed the transcription level of every gene in the three 601 strains and in the wild-type strain. We compared the expression level in the 601 array with the rest of the genes by ordering every transcript according to its counts-per-million (cpm) value. Respectively 4.28, 1.94 and 12.7 per million transcripts were identified in the 167, 197 and 237 bp NRL strains. When ordered from high to low expression, transcripts containing the 601 sequence ranked respectively at position 5325, 5522 and 4789, meaning that in each strain more than 80% of the genes expressed are detected at a higher level. If we take into consideration the repeated nature of the array, the number of transcripts should also be divided by the approximate number of repeat within each array (∼50), leading to an even weaker transcription activity of the 601 arrays (Table 1).

**Table 1.** 601 repeats inserted at the YMR262 locus are weakly transcribed.

| 601 array type | cpm | rank / total genes<br>which cpm is > 0 | rank 601 monomer/ total genes<br>which cpm is > 0 |
| --- | --- | --- | --- |
| 167 | 4.28 | 5325 / 6068 | 5915 / 6068 |
| 197 | 1.94 | 5522 / 6031 | 5928 / 6031 |
| 237 | 12.7 | 4789 / 6014 | 5711 / 6014 |

## Discussion

The in vivo spacing between nucleosomes has been proposed to play an important role in many chromosome functions ranging from the definition of chromatin states and gene activity [23] to homologous pairing [24]. In this study we built synthetic DNA arrays in the genome of *S. cerevisiae*, as a first step to address experimentally the biological effects of unnatural nucleosome spacing. Based on the extensive published work on nucleosome positioning in vitro, we chose the 601 nucleosome-positioning sequence to assemble repeats of the artificial 601 sequence according to three theoretical NRLs (167, 197 and 237 bp). First we showed that we can assemble over 15 kb long arrays of the 601 sequence, as we previously did with other repeat sequences [21]. While we confirm that regular 167, 197 and 237 NRL nucleosomal arrays can be formed using the 601 sequence in vitro, the results reported here reveal that the 601 sequence is not able to position nucleosomes in vivo. We show unambiguously that synthetic arrays of 601 DNA in vivo does not lead to the formation of a synthetic chromatin fragment with controlled nucleosome positions. Nonetheless, nucleosomes are not randomly organized on these synthetic arrays (Figure 4A,B and C) and tend to be positioned away from the 601 sequence.

Taken together, our results can be explained as follows. Firstly, the DNA sequence motifs that would position nucleosomes are different in vivo and in vitro. This result is in line with our recent deep learning based mutational screen of the *S*.*Cerevisiae* genome which did not point at any DNA motifs that would attract and position nucleosomes in vivo [25]. Secondly, the main driver of nucleosome spacing is not the DNA sequence but the yeast chromatin folding machinery, which yields a consistent spacing of 163 bp (Figure 3 B). On our synthetic arrays, the observed spacing is slightly larger, ranging from 166 to 183 (Figure 3 D) but is still much smaller than the spacing that would be dictated by the DNA sequence alone for the 197 and 237 bp arrays. Based on these two principles, we can explain that in the case of the 167 bp array, since the periodicity of the sequence is close to the natural periodicity of nucleosome positioning in yeast, most nucleosomes are phased with the sequence and we observe a preferential positioning of the nucleosome centers on the monomer sequence (Figure 5 top). For the 197 bp constructs on the other hand, the spacing imposed by the yeast molecular machinery in not in phase with the DNA array and we observe a more uniform distribution of nucleosomal centers, although there is still a slight preference for positioning outside of the 601 core (Figure 5, middle). Finally, in the case of the 237 bp repeat, the impose spacing is such that 3 nucleosomes would occupy approximately 2 DNA repeats, resulting in three fuzzy peaks in the distribution of the nucleosome centers (Figure 5, bottom). In conclusion, this study emphasizes that the 601 sequence should not be assumed to maintain nucleosome positioning when inserted in genomic loci in vivo, and new approaches will be needed to design in vivo nucleosome positioning sequences for the engineering of designer chromatin.

**Figure 5.**
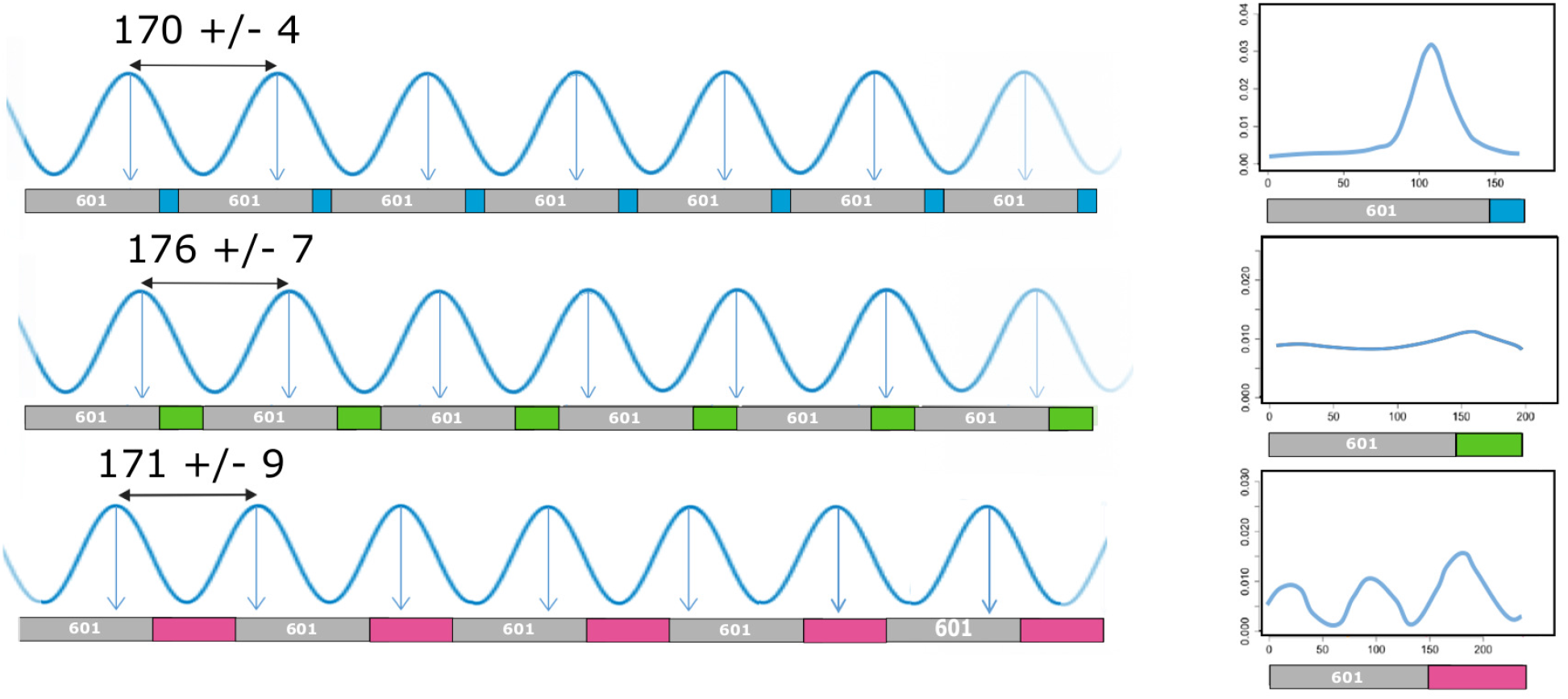
A model of nucleosome positioning on each of the 601 arrays in vivo. Proposition of nucleosome organization over 5-8 repeats (left panels based on both our NRL measures on gel and on the experimental distributions of nucleosomal fragment midpoints on each 601 repeats (right panels).

## Materials and Methods

### Strains, plasmids, reagents and media

The VL6-48 strain (ATCC MYA3666) was used for the in vivo assembly of DNA repeats into yeast 2 micron plasmids [26]. In vivo genomic assembly of 601 DNA repeats was achieved in the yeast strain YPH499 [27]. Strains derived from this study are listed in Supplementary Table S1. Plasmids used for the in vivo expression of spCas9 and the guide RNA targeting the YMR262 gene have been previously described [21] and are listed in Supplementary Table S2. Strains were grown at 30°C in yeast extract Peptone Adenine Dextrose 2% media (YPAD) or in the appropriate synthetic complete Dextrose 2% media (SCD) media minus relevant amino acids necessary to maintain plasmid borne auxotrophic markers. All media reagents were purchased from Formedium and used as recommended. Oligonucleotides used in this study were synthetized by Eurogentec. Enzymes for nucleic acids modification were purchased from New England Biolabs. Zymolyase 20T was purchased from Amsbio.

### Assembly of 601 tandem DNA repeats in the yeast genome

Assembly of 601 DNA repeats on an episomal vector in vivo was achieved using a method adapted from the Transformation Associated Recombination method [28]. The full method can be found in the Supplementary Material and Methods. Direct in vivo assembly using CRISPR/Cas9 and overlapping oligonucleotides in the Chromosome XIII of *S. cerevisiae* strain YPH499 was performed as previously described [21]. Briefly, donor DNA containing the left (YMR/601) and the right (601/YMR) genomic junction were amplified by PCR using primer couples AL-O-14/15 and AL-O-16/17 respectively. These donor DNA results, upon recombinational assembly, in the deletion of the region -129 to 232 bp of the YMR gene. All oligonucleotides used for in vivo repeat assembly are listed in Supplementary Table S3. YPH499 was first transformed with the CAS9 expressing plasmid p414-TEF1p-Cas9-CYC1t. This plasmid was a gift from George Church (Addgene plasmid *#* 43802) [29]. The resulting strain was transformed using the LiAc technique [30] with 1 *µ*g of gRNA expressing plasmid targeting YMR262 (pAL31), 100 *pmol* of each of the four or six appropriate 601-oligonucleotides (all oligonucleotides and plasmids are listed in supplementary Table S2 and S3), and 10 *pmol* of both YMR/601 and 601/YMR junction PCR. After transformation cells were plated on SCD-His-Ura to select for cells carrying both CAS9 and gRNA expressing plasmids. To test for correct in vivo assembly of 601-tandem repeated arrays, each genomic junction between the assembled repeated array and genomic DNA were analyzed by Sanger sequencing with primers AL-O-02/22 and AL-O-01/23, to amplify respectively the left YMR/601 junction or the right 601/YMR junction (see Supplementary Figure S2). To confirm assembly at the correct locus and to estimate the size of the assembled arrays, recombinant clones were also analyzed by southern-blotting. Genomic DNA from recombinant strains were digested with BamhI and DraI, which cut at each side of the insertion locus. Digested DNA was electrophoresed in 1% agarose and transferred by capillarity onto a nylon membrane (Hybond N+, GE healthcare). Membranes were hybridized in Church buffer at 68°C with two different ^32^P-radiolabeled probes (Church and Gilbert, 1984). The 601 probe was made of the 147 *bp* long 601-core sequence. The genomic probe was a 1 *kb* DNA fragment amplified by PCR from genomic DNA using primers AL-O-24/25 (see Table S3). Probes were labeled with the prime-it II random primer labelling kit (Agilent) using *α*^32*P*^ -dCTP. For each 167, 197 and 237 design the clone carrying the longest 601 tandem repeats was named respectively ALY1, ALY2 and ALY3 and selected for further analysis (Figure 2). Membranes were scanned using a FLA 9500 GE healthcare Scanner at 200 *µ*m resolution.

### Chromatin digestion by Micrococcocal Nuclease (MNase)

Each strain was grown to an OD_600_ of 0.8 in 250 *mL* Synthetic Complete Dextrose 2 % media at 30°*C* with shaking at 200 *rpm*. Cultures were treated with a final concentration of 1.85 % formaldehyde for 30 min at 30°C. Cross-linking was stopped by addition of 105 *mM* Glycine (final concentration) and collected by centrifugation (6500*g*, 10 *min*). Cell pellets were washed and resuspended in 50 *mL* of 1 *M* Sorbitol, 10 *mM* Tris *pH* 7.5 supplemented with 10 *mM β*-mercaptoethanol and 15 *mg* of Zymolyase 20T. Cells were further incubated at 30°C with 1h shaking at 100 *rpm*. Spheroplasts were pelleted (6500*g*, 10 *min*), and resuspended in 2.4 *mL* solution containing 1*M* Sorbitol, 50 *mM* NaCl, 10 *mM* Tris *pH* 7.5, 5 *mM* MgCl_2_, 1 *mM* CaCl_2_ and 0.75 % Igepal Ca630 freshly supplemented with 1 *mM β*-mercaptoethanol, 500 *µM* spermidine and various amounts of MNase (between 6000 and 30000 units), to allow choosing samples with the appropriate digestion pattern for further processing. The spheroplasts/MNase mixture was then incubated at 37°C for precisely 30 *min*. The reaction was stopped by adding 600 *µL* of 1% SDS, 10 *mM* EDTA. Reversal of crosslink and protein removal was achieved by adding 0.6 *mg* of Proteinase K (Invitrogen) and overnight incubation at 65°C. Samples were extracted with phenol/chloroform, and DNA was ethanol precipitated treated with DNase-free RNase.

### MNase-gel analysis

The MNase-gel analysis was performed as follows. Mildly digested Mnase resistant nucleosomal DNA was separated by electrophoresis in a 1.3 % agarose gel. After migration, the gel was colored with Ethidium Bromide (0.5*µg*/mL) and imaged in a Gbox bioImager (Syngene UK). The resulting image was analysed using a simple image analysis routine developed in-house in Matlab to calculate the size of digested nucleosomal DNA relative to Molecular Weight standards (1 kb+ DNA (Thermo Fischer Scientific)). Following gel imaging, the DNA was transferred on a nitrocellulose membrane by capillarity using the Southern blot protocol described above. Membranes were hybridized with a 147 bp *α*^32*P*^ -labelled probed and exposed against a phosphor plate for 24-48 hours. Phosphor plates were scanned using a Typhoon FLA 9500 scanner at 200 *µ*m resolution (GE Healthcare). The image obtained was analyzed with the same image analysis routines applied to gels colored with Ethidium Bromide.

### Preparation of DNA libraries for MNase-seq analysis

For the preparation of fragment libraries, 500 *ng* of gel purified mononucleosomal DNA were repaired using the PreCR Repair Mix Kit (New England Biolabs) with 100 *µM* dNTPs in a 50 *µL* reaction, as recommended by the manufacturer. The reaction was incubated for 30 minutes at 37°C, purified using the QIAquick PCR Purification Kit (Qiagen) and eluted in 80 *µL* H_2_O. Nucleosomal DNA was 5’ phosphorylated by adding 333 *µM* dNTPs, 50 units of T4 polynucleotide kinase (NEB), 15 units of T4 DNA polymerase (NEB) and 5 units of Klenow DNA polymerase (NEB) to 80 *µ*L of repaired mononucleosomal DNA to a final reaction volume of 120 *µ*L. The reaction was incubated 30 minutes at room temperature, purified using the QIAquick PCR Purification Kit and eluted in 30 *µ*L H2O. To add a 3’-dA to the nucleosomal DNA the 30 *µ*L eluate was completed to 50 *µ*L with 0.2 mM dATP and 15 units Klenow DNA polymerase (3’-5’ exo-). The reaction was incubated 30 minutes at 37°C and inactivated 10 minutes at 65°C. Reaction was purified using the QIAquick PCR Purification Kit, and eluted in 20 *µ*L. Adapters were then ligated to the nucleosomal DNA by completing the reaction to 30 *µ*L with 10 mM adapters and 1200 units of T4 DNA ligase. The reaction was incubated at 16°C overnight, followed by inactivation for 20 minutes at 65°C. Ligated DNA was purified by gel electrophoresis. The adapter-ligated nucleosomal DNA was amplified by PCR prior to sequencing, using Illumina PE1.0 and PE2.0 primers. We used only 8 to 15 cycles to minimize possible bias due to PCR amplification. PCR were as follows: 3 *µ*L of adapter-ligated nucleosomal DNA were used in a 50 *µ*L reaction volume containing 5 *µ*L of each Illumina paired-end PCR primers at 2 *µ*M, 1 *µ*L of 10mM dNTPmix and 2 *µ*L of Phusion polymerase (NEB). PCR products were purified using the QIAquick PCR purification column and eluted with 30 *µ*L Qiagen Elution Buffer. DNA concentration was determined using a Qubit Fluorometer. Libraries were multiplexed and then sequenced on a HiSeq 2000 device.

### MNase-seq analysis

For each 601-containing strains four individual libraries were sequenced, yielding 5 to 28 Million reads for each library. For the three 601 designs we constructed a reference genome carrying two full repeats plus 65 bp of a third one. Indeed, the construction of the chimeric genome is required to take into account all possible alignments of the reads. Reads for which one mate overlaps the linker and the beginning of the following 601-core will align on a second repeat in our reference genome. Moreover, extreme cases where the 2 mates of a read overlap two linkers will align on the beginning of a third repeat (65 bp were chosen as the length of our read -1 bp) (see Supplementary Figure S1). After removing of barcodes, paired-end reads of 65 nt each end were mapped against the appropriate 601 (167, 197 or 237) reference genome using Bowtie2 (version 2.3.5.1) [31, 32] allowing at most 2 alignments per read (-k 2) and fragments length ranging from 100 to 200 bp for 167 and 197 strains, and from 120 to 250 bp for 237 strains (-I and -X parameters). By choosing a reference genome with two repeats and 65 bp of an additional 601, a large proportion of the reads of interest should be able to map twice. For this reason we allowed 2 hits per read using Bowtie2. Concordant reads were specifically selected using samtools (version 1.10). Reads presenting multiple hits on the repeat region were filtered to select the most 5′ alignment using a python script. Coverage per base-pair was determined in both the whole-genome (WG) and the 601-repeats in each individual library, to calculate the enrichment factor in the repeats (see Supplementary Figure S6). For all the fragments overlapping the 601 area their midpoint positions were localized on the repeats using a Python script. Finally, these midpoint coordinates were merged on a unique repeat to represent the averaged nucleosomal dyad density at each position of a single repeat using a custom R script.

### RNA libraries preparation

Total RNA was extracted by starting from 25 mL culture of each strain grown to an OD_600_ of 0.5 in SCD media at 30°C with shaking at 200 rpm. Cells pellets were resuspended in 500 *µ*L of Nucleazol solution (Macherey Nagel, 740404.200) followed by 20 minutes agitation in MN Bead Tubes Type C (Macherey Nagel, 740813.50). Total RNA was then purified using the NucleoSpin® RNA Set for NucleoZOL (Macherey Nagel, 740406). RNA concentration was determined using a ND-1000 nanodrop spectophotometer. Following a rRNA depletion step, purified RNA was fragmented and reverse-transcribed into 300 bp cDNA before single-end Illumina adaptor ligation. Finally, libraries were sequenced on a HiSeq 4000 device by the Eurofins sequencing platform (INVIEW Transcriptome Explore).

### RNA-seq analysis

We sequenced RNA from the three 601-strain and the wild-type stain during exponential growth phase in YPAD media. 50 nt long reads were pseudoaligned to the reference protein-coding cDNA collection of the *S. cerevisiae* strain S288C via Kallisto [33] (version 0.46.2) in the single-end mode. The YMR262 cDNA sequence of the fasta files used to create index was replaced by the sequence of four 601 repeats of the appropriate design when aligning the three 601 strains. For each of the four biological replicates cpm (counts-per-million) values for every gene were calculated with the edgeR (version 3.30.3) package of Bioconductor [34]. cpm values were normalized using edgeR to correct for sample-specific variation typically introduced by differences in library size. To verify that 601 repeats do not disturb the global transcriptome we calculated the *log*2(*cpm*) value of every gene (except the ones for which all samples displayed less than 1 *cpm*) to determine expression fold change between the wild-type strain and the three 601 strains (see Supplementary RNAseq.csv file). To estimate the 601 transcription rate we deduced in each 601-strain from the cpm value in the repeats the rank of expression of the 601 region over the 6612 genes analyzed (see Table 1).

## Acknowledgments

The authors thank the members of the Genome Structure and Instability unit for discussions and Dr Romain Koszul for providing assistance with sequencing. This work was supported by a grant from the national research agency (MuSeq, ANR-15-CE12-0015) and by core funding from CNRS, INSERM and MNHN to the Structure and Genome instability unit. French Museum of National History to the Genome Structure and Instability unit. AL was supported by a doctoral fellowship from the National Museum of Natural History. Work in the group of VC is part of “Institut Pierre-Gilles de Gennes” (“Investissements d’Avenir” program ANR-10-IDEX-0001-02 PSL and ANR-10-LABX-31) and the Qlife Institute of Convergence (PSL Univesité).

## Appendix A. Supplementary Data

Supplementary Methods and data available online at XXX.

## Conflict of interest statement

V.C. is cofounder of PicoTwist.

## data availability

Sequencing data are available for review here:

