## supplemental methods and figures for "Nucleosome positioning on large tandem DNA repeats of the ‘601’ sequence engineered in *Saccharomyces cerevisiae*"

Lancrey, A & al.

### Supplementary Material & Methods

**Genomic stability of 601-DNA tandem repeats upon mitotic propagation**

Yeast clones were grown in 5ml YPD overnight and plated on YPD agar. 96 colonies were picked for each plate and grown in 0.5 ml YPD. After overnight growth, the 96 cultures for each repeat size were divided in 8 pools of 12 cultures and processed by preparation of genomic DNA. Genomic DNA pools were digested by BamHI and DraI. Separation of digested DNA and Southern blotting was achieved as described in the main Material and Methods section.

**Engineering of plasmids carrying tandem DNA repeats of the 601 sequence**

601 oligonucleotides were assembled *in vivo* into a plasmid by a procedure derived from the Transformation-Associated recombination (TAR) method [1] using the VL6-48 strain (ATCC # MYA-3666). VL6-48 was transformed simultaneously by both the linearized "host plasmid" and the indicated overlapping oligonucleotides. The "host plasmid" used is derived from pRS423 [2], which Multi Cloning Site was previously modified so that restriction digestion with ZraI restriction enzyme results in a linearized plasmid with 5' and 3' ends being homologous to the ends of the repeats over 34 and 42 nucleotides respectively. This plasmid is named pRS423-NS-601. $10^8$ spheroplasts were transformed with a DNA mixture containing 40 fmol of the pRS423-NS601 plasmid linearized with ZraI and 4 pmol of each of the four or six indicated repeat-coding oligonucleotides, the volume of the DNA mixture not exceeding 40 µL. After transformation spheroplasts were mixed with melted 3%-agar-SCD-His-1M SORBITOL and plated on SCD-His-1M SORBITOL agar plates to select for cells carrying the pRS423-NS-601 plasmid. Grown transformants were further screened for plasmids carrying 601 repeats by yeast colony hybridization [3]. Reisolated colonies were directly lifted onto a nylon membrane and digested with β- glucuronidase. Cells were lysed in a NaOH containing buffer to release DNA onto the membrane. Membranes were incubated with a radiolabeled DNA probe carrying the 601 sequence overnight and the radioactive signal was analyzed by scanning on a Storm Phosphorimager (GE). Positive colonies giving a strong radioactive signal were selected for further analysis. Plasmids carried by the positive clones were extracted by yeast plasmid miniprep purification and transformed into the E. coli XL10 strain. Recombinant plasmids were purified and analyzed by gel electrophoresis after NotI/SpeI restriction digest. For each of the three 601 designs, the largest repeats were 3.3 kb, 2.7 kb and 3.3 kb long for respectively pRS-601-167, pRS-601-197 and pRS-601-237. These three plasmids were used for *in vitro* chromatin reconstitutions and their 601 insert was further characterized by single molecule fingerprinting.

**Single molecule fingerprinting**

Plasmids containing 601 repeats were linearized by NotI and SpeI restriction enzymes. Reaction was resolved by electrophoresis and gel purified. 3 pmol of the repeated DNA array was then ligated using T4 DNA ligase (NEB) to a hairpin forming oligonucleotide on the NotI side and to a Y-shape DNA on the SpeI side, according to [4]. After gel purification, the substrate was finalized with addition of 2.5 mM dA/dC (Invitrogen), 1mM dig-dUTP (Roche) and 20 units Klenow polymerase. This last step allows addition of digoxygenin moieties in the double-stranded handle of the molecule and serves to the attachment of the molecule to the anti-dig coated coverslip. Attachment of the molecule to beads, loading the flow cell and mechanical sequencing was achieved as described by Ding et al [5]. Fingerprinting results are displayed Figure S4. The number of repeats present in each molecule was detected by the number of pauses observed during open/close cycles, each pause corresponding to a binding event of the Oligo which sequence is displayed for each design. According to the observed pause events the 601-167, -197 and -237inserts carried respectively 20, 14 and 14 repeats. This corresponds to inserts of 3340 bp for the 167 bp monomer, 2728 bp for the 197 bp monomer and 3275 bp for the 237 monomer. These results are compatible with sizes calculated from results of gel electrophoresis of the inserts.

**In vitro chromatin reconstitution**

Salt jump dialysis was used to assemble nucleosome arrays on the three pRS-167, pRS-197 and pRS-237 plasmids. 1.6µg of each DNA were incubated in 5.3 µL reactions containing 2M NaCl and 1.8 µg of HeLa core histones (Active Motif # 53501). The DNA/histones mixtures were incubated at 37°C for 10 minutes. 3 volumes of BSA (100µg/ml in TE Buffer) were then added to the samples followed by incubation at 37°C for 30 minutes. The samples were then dialyzed for two hours at room temperature by placing them in a microdialysis tubing with the dialysis membrane contacting Tris-EDTA buffer. Samples were then treated with 1.5 units of MNase for 10 minutes at room temperature. Digestion was stopped after 10 min by adding 25 µL of 0.5M EDTA. Samples were then treated with 25 µg of proteinase K for 30 min at 50◦C. Samples were purified with MinElute PCR Purification Kit (Qiagen 28004) then verified on a 1%agarose gel. To obtain the three 601-167, -197 and -237 ladders each naked plasmid was digested with excess of XbaI restriction enzyme for 2 hours, which cut once per repeat (Figure S1). The obtained products were migrated on an electrophoresis gel to purify the band containing all the monomer repeats. Once purified 600 ng of the monomers were submitted to a partial ligation with 1200 units of T4 DNA ligase 10 minutes at room temperature followed by inactivation 10 minutes at 65°C to reconstitute the corresponding 601 ladder. Southern blot analysis of the digested reconstitutions were performed with the 601 radiolabeled probe made from a 147 bp long 601 core sequence following the same protocol than described in the material section of the main text.

#### A. 167 bp repeats – 601-167 (2 monomers + 601 core)

f1-167

ccgttgctgcgtgctgaaggggattaattaatACAGGATGTATATATCTGACACGTGCCTGGAGACTAGGGAGTA

f3-167

ATCCCCTTGGCGGTTAAAACGCGGGGGACAGCGCGTACGTGCGTTTAAGCGGTGCTAGAGCTGTCTACGACCAAT

r2-167

TGAGCGGCCTCGGCACCGGGATTCTCCAGGGCGACCGCGTCTAGAGTCCACAGGATGTATATATCTGACACGTGC

r4-167

CTGGAGACTAGGGAGTAATCCCCTTGGCGGTTAAAACGCGGGGGACAGCGCGTACGTGCGTTTAAGCGGTGCTAG

AGCTGTCTACGACCAATTGAGCGGCCTCGGCACCGGGATTCTCCAGGGCGACCGCGTCTAGAGTCCACAGGATGT

ATATATCTGACACGTGCCTGGAGACTAGGGAGTAATCCCCTTGGCGGTTAAAACGCGGGGGACAGCGCGTACGTG

CGTTTAAGCGGTGCTAGAGCTGTCTACGACCAATTGAGCGGCCTCGGCACCGGGATTCTCCAGacaaggtttctc

actatcaagatgtactggagtataagaatg

#### B. 197 bp repeats – 601-197 (2 monomers + 601 core)

f1-197

ccgttgctgcgtgctgaaggggattaattaatACAGGATGTATATATCTGACACGTGCCTGGAGACTAGGGAGTA

f3-197

ATCCCCTTGGCGGTTAAAACGCGGGGGACAGCGCGTACGTGCGTTTAAGCGGTGCTAGAGCTGTCTACGACCAAT

r2-197 f5-197

TGAGCGGCCTCGGCACCGGGATTCTCCAGGGCGACCGCGTCTAGAGTCCATCACATAAGGGATGAACTCGGTGTA

r4-197

GAACACAGGATGTATATATCTGACACGTGCCTGGAGACTAGGGAGTAATCCCCTTGGCGGTTAAAACGCGGGGGA

r6-197

CAGCGCGTACGTGCGTTTAAGCGGTGCTAGAGCTGTCTACGACCAATTGAGCGGCCTCGGCACCGGGATTCTCCA

GGGCGACCGCGTCTAGAGTCCATCACATAAGGGATGAACTCGGTGTAGAACACAGGATGTATATATCTGACACGT

GCCTGGAGACTAGGGAGTAATCCCCTTGGCGGTTAAAACGCGGGGGACAGCGCGTACGTGCGTTTAAGCGGTGCT

AGAGCTGTCTACGACCAATTGAGCGGCCTCGGCACCGGGATTCTCCAGacaaggtttctcactatcaagatgtac

tggagtataagaatg

#### C. 237 bp repeats – 601-237 (2 monomers + 601 core)

f1-237

ccgttgctgcgtgctgaaggggattaattaatACAGGATGTATATATCTGACACGTGCCTGGAGACTAGGGAGTA

f3-237

ATCCCCTTGGCGGTTAAAACGCGGGGGACAGCGCGTACGTGCGTTTAAGCGGTGCTAGAGCTGTCTACGACCAAT

r2-237 f5-237

TGAGCGGCCTCGGCACCGGGATTCTCCAGGGCGACCGCGTATAGGGTCCATCACATAAGGGATGAACTCGGTCTA

r4-237

GAACTATCCGACTGGCACCGGCAAGGTCGCTGTTCAATACATGCACAGGATGTATATATCTGACACGTGCCTGGA

r6-237

GACTAGGGAGTAATCCCCTTGGCGGTTAAAACGCGGGGGACAGCGCGTACGTGCGTTTAAGCGGTGCTAGAGCTG

TCTACGACCAATTGAGCGGCCTCGGCACCGGGATTCTCCAGGGCGACCGCGTATAGGGTCCATCACATAAGGGAT

GAACTCGGTCTAGAACTATCCGACTGGCACCGGCAAGGTCGCTGTTCAATACATGCACAGGATGTATATATCTGA

CACGTGCCTGGAGACTAGGGAGTAATCCCCTTGGCGGTTAAAACGCGGGGGACAGCGCGTACGTGCGTTTAAGCG

GTGCTAGAGCTGTCTACGACCAATTGAGCGGCCTCGGCACCGGGATTCTCCAGacaaggtttctcactatcaaga

tgtactggagtataagaatg

**Figure S1 : Sequence of the designed repeats showing 2 monomers (167 (A), 197 (B) and 237 (C) bp) and a core 601 sequence in the genomic context of chromosome XIII.** Inserted monomers are shown in capital letters, genomic DNA in small letters. 601 core sequence is highlighted in red, linker sequences are highlighted in yellow, Xba site is underlined. Position of the overlapping oligonucleotides (Table S3) are indicated for each design.

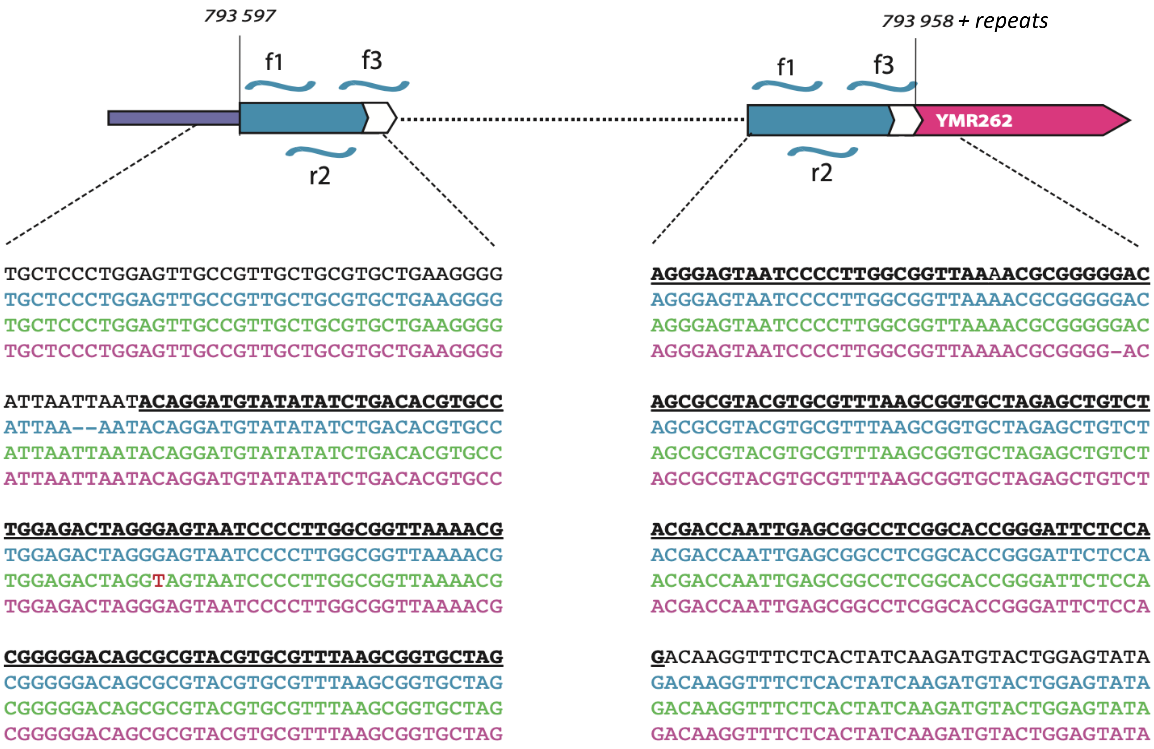


**Figure S2 : Analysis of the genomic DNA-601 junctions by Sanger sequencing**. Sequencing results are shown for the three 601 repeats with the blue lagoon, pink and green sequences corresponding respectively to the 167, 197 and 237 bp repeats. The left and right panels correspond respectively to the left YMR262/601 and right 601/YMR262 junction sequences. The black sequence corresponds to the theoretical sequence, the nucleotides belonging to the 601 sequence are underlined. Substitutions are indicated by red letters and deletions by indels. Sequencing covers the beginning of the 1st 601 repeat until the end of f3 on the left side and the last repeat from the middle of f1 on the right side.

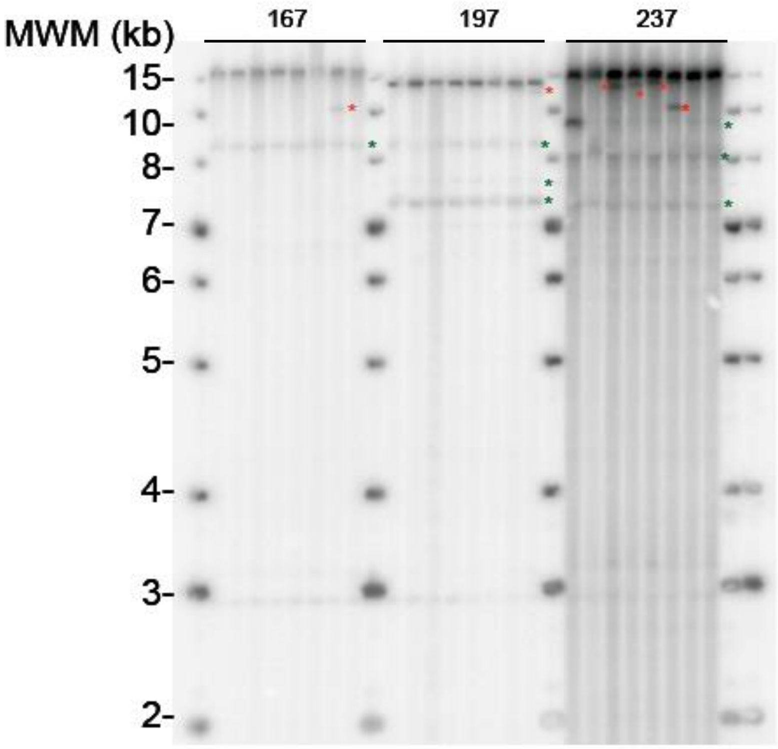


**Figure S3: Genomic stability of 601 tandem DNA repeats in vivo.** Southern blot analysis of genomic DNA prepared from three propagated clones containing the 601 tandem DNA repeats with 167, 197 and 237 bp monomers respectively. Cells were propagated for ~30 divisions before harvesting for DNA preparation (see Supplementary Material and Methods). Each lane contains 12 pooled cultures processed together. 96 cultures were therefore processed for each propagated clone. Red stars denote size variant, green stars denote variants present in all the clone and were therefore present in the original culture. They were not taken into consideration in the estimation of genomic instability.

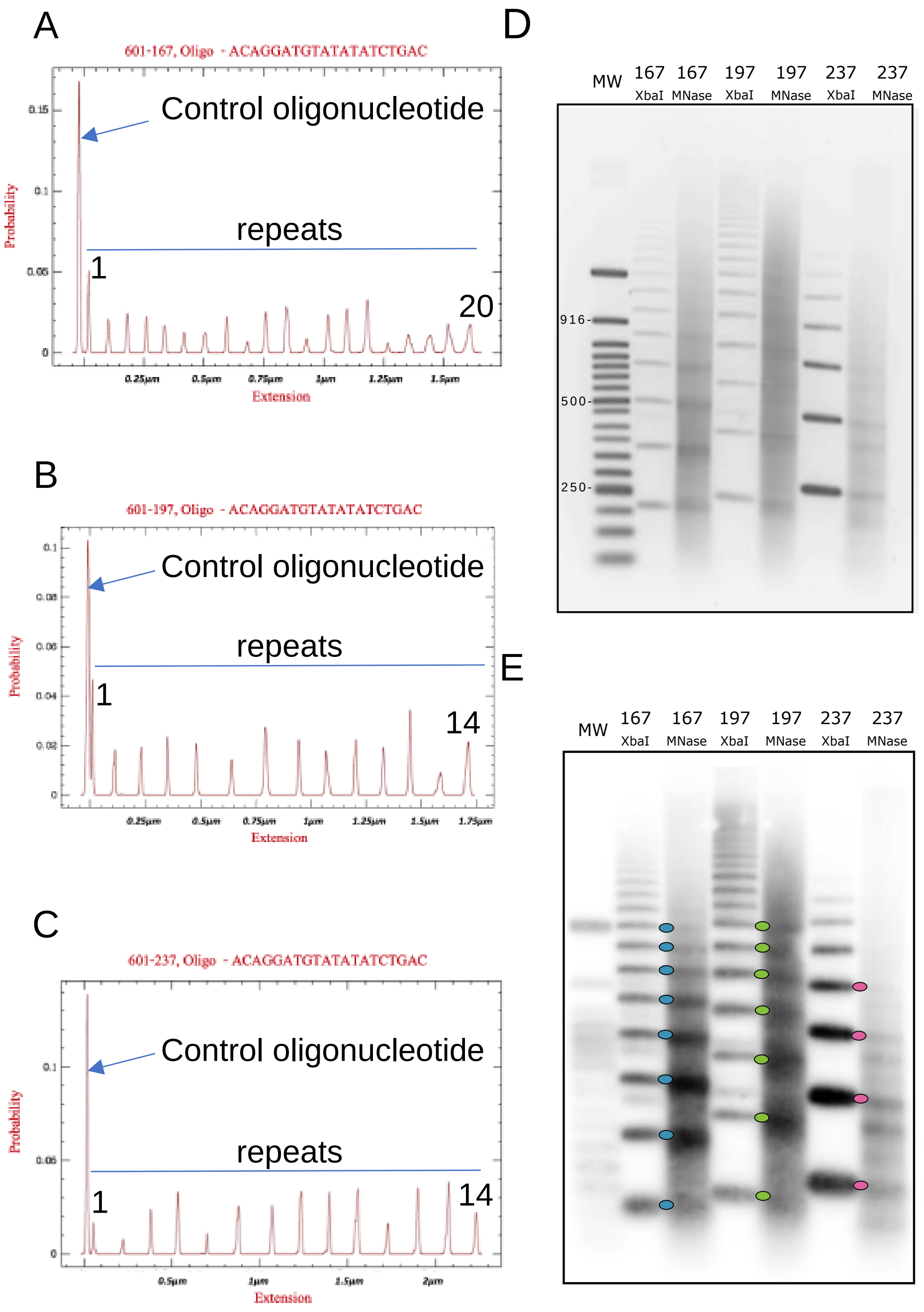


**Figure S4 : Characterization of 601 nucleosomes assembled in vitro on plasmid DNA. (A-C)** Single molecule fingerprinting of the engineered plasmids. The binding probability of the indicated 601-oligonucleotide was calculated along the rezipping time course of the total DNA molecule carrying the repeats for respectively the 167 **(A)**, 197 **(B)** and 601-237 repeats **(C)** contained in the respective plasmid. **(D, E)** In vitro reconstitution of chromatin. D. EtBr gel showing the Mnase digested chromatin (Mnase) versus DNA ladder of respectively 167, 197 and 237 bp steps (XbaI lane). **(D)** The gel was then transferred onto a membrane by Southern blotting and hybridized with a 601 radiolabeled probe to determine the NRL in the 601repeats of each of the three chromatinized plasmids. The colored dot were drawn to relate the ladder band to its corresponding nucleosomal band in the ”MNase” lane.

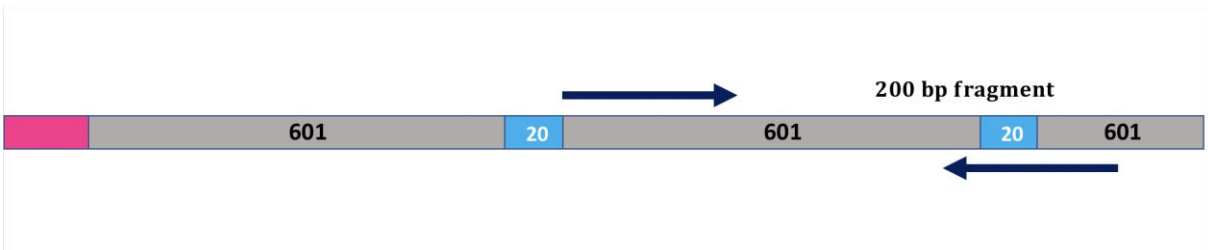


**Figure S5 : Structure of 601-167 repeats in the reference genome.** The grey rectangles correspond to the 601 147bp core DNA sequence, the blue rectangle to the 20 bp DNA linker and the pink rectangle to the left genomic junction. Arrows represent concordant mates that allow a nucleosomal DNA fragment to be reconstructed. Here is represented an example of a fragment which could not be considered with a smaller 601 repeated area. In this case the first read overlaps two 601-167 repeats so that it can not map on the first repeat. With this 200 bp long fragment the second read also overlaps two repeats. In this case the first read overlaps only one bp of the first repeat the second read overlaps 32 bp of a third repeat.

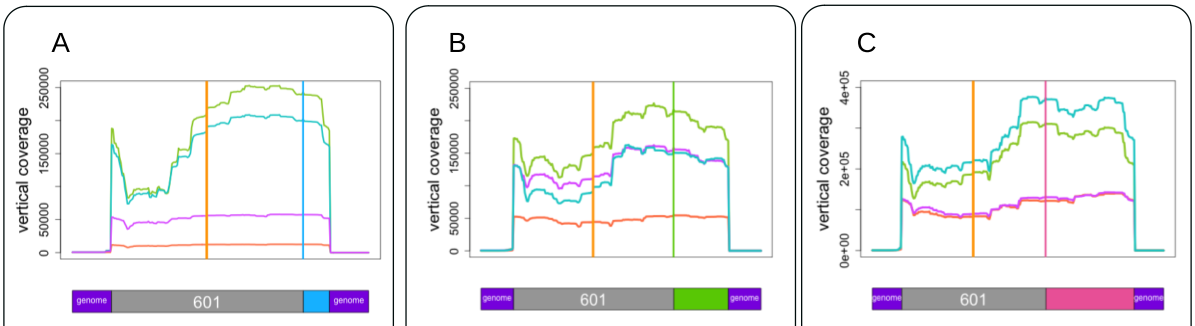


D

| Repeats | WG | |
|---|---|---|
| coverage/bp | coverage/bp | Enrichment |
| 4 664 | 63 | 74 |
| 21 707 | 310 | 70 |
| 80 023 | 283 | 283 |
| 97 489 | 250 | 389 |

E

| Repeats | WG | |
|---|---|---|
| coverage/bp | coverage/bp | Enrichment |
| 18 735 | 57 | 330 |
| 60 927 | 309 | 197 |
| 62 886 | 211 | 298 |
| 85 576 | 351 | 244 |

F

| Repeats | WG | |
|---|---|---|
| coverage/bp | coverage/bp | Enrichment |
| 46 892 | 65 | 717 |
| 49 542 | 287 | 172 |
| 135 315 | 220 | 616 |
| 110 620 | 185 | 598 |

**Figure S6 : Comparison of vertical coverage in the Whole Genome versus the 601-repeats.** The same color of each individual library of each strain is the same as in Figure 4. Vertical coverage was plotted on a unique repeat with its genomic boundaries over 30 bp in the 601-167 strain **(A),** in the 601-197 strain **(B)** and in the 601-237 strain **(C).** Vertical coverage per bp was calculated independently for the Whole Genome (WG) and the 601-repeats and enrichment factors in the repeats were deduced in the 601-167 strain **(D),** in the 601-197 strain **(E)** and in the 601-237 strain **(F).**

Table S1. Strains used in this study.

| Strain name | Genotype | Reference |
|---|---|---|
| VL648 | MATα, his3- Δ200, trp1-Δ1, ura3–52, lys2, ade2–101, met14 | SGD |
| YPH499 | MATa ura3-52 lys2-801_amber ade2-101_ochre trp1-Δ63 his3-Δ200 leu2-Δ1 | SGD |
| ALY0 | YPH499 + pAL30 | *Lancrey & al. Sci. Rep. 2018* |
| ALY1 | ALY0 ymr262::601-167 | This study |
| ALY2 | ALY0 ymr262::601-197 | This study |
| ALY3 | ALY0 ymr262::601-237 | This study |

Table S2. Plasmids used in this study.

| **Plasmid Name** | **Description** | **Reference** | **plasmids/oligos** |
|---|---|---|---|
| pRS-601 | pRS423-601-hooks | This study | pRS423 |
| pRS-601-167 | pRS423-601-167-repeats | This study | pRS-601/ AL-O-01 - 04 |
| pRS-601-197 | pRS423-601-197-repeats | This study | pRS-601/ AL-O-08 -13 |
| pRS-601-237 | pRS423-601-237-repeats | This study | pRS-601/ AL-O-01 - 03 / AL-O-05 - 07 |
| p414-Cas9 | p414-TEF1p-Cas9-CYC1t | Addgene | - |
| pAL30 | pRS413-Cas9 | *Lancrey & al. Sci. Rep. 2018* | pRS413 ; p414 |
| p426 | p426-SNR52p-gRNA.CAN1.Y-SUP34t | Addgene | - |
| pAL31 | p426-SNR52p-gRNA.YMR262-SUP34t | *Lancrey & al. Sci. Rep. 2018* | p426 / AL-O-18/19; AL-O-20/21 |

Table S3. Oligonucleotides used in this study.

| **Primer Name** | **Sequence** | **Description** |
|---|---|---|
| AL-O-01 | ACAGGATGTATATATCTGACACGTGCCTGGAGACTAGGGAGTAATCCCCTTGGCGGTTAA | f1-167/237 |
| AL-O-02 | CTAGCACCGCTTAAACGCACGTACGCGCTGTCCCCCGCGTTTTAACCGCCAAGGGGATTAC | r2-167/237 |
| AL-O-03 | CGTGCGTTTAAGCGGTGCTAGAGCTGTCTACGACCAATTGAGCGGCCTCGGCACCGGGAT | f3-167/237 |

| | | |
|---|---|---|
| AL-O-04 | GTGTCAGATATATACATCCTGTGGACTCTAGACGCGGTCGCCCTGGAGAATCCCGGTGCCGAGG | r4-167 |
| AL-O-05 | CCGAGTTCATCCCTTATGTGATGGACCCTATACGCGGTCGCCCTGGAGAATCCCGGTGCCGAGG | r4-237 |
| AL-O-06 | CATCACATAAGGGATGAACTCGGTCTAGAACTATCCGACTGGCACCGGCAAGGTCGCTG | f5-237 |
| AL-O-07 | CTCCAGGCACGTGTCAGATATATACATCCTGTGCATGTATTGAACAGCGACCTTGCCGGTGCCAGTC | r6-237 |
| AL-O-08 | ACAGGATGTATATATCTGACACGTGCCTGGAGACTAGGGAGTAATCCCCTTGGCGG | f1-197 |
| AL-O-09 | TAAACGCACGTACGCGCTGTCCCCCGCGTTTTAACCGCCAAGGGGATTACTCCC | r2-197 |
| AL-O-10 | CGGGGGACAGCGCGTACGTGCGTTTAAGCGGTGCTAGAGCTGTCTACGACCAATTG | f3-197 |
| AL-O-11 | CGGTCGCCCTGGAGAATCCCGGTGCCGAGGCCGCTCAATTGGTCGTAGACAGCTC | r4-197 |
| AL-O-12 | GCACCGGGATTCTCCAGGGCGACCGCGTCTAGAGTCCATCACATAAGGGATG | f5-197 |
| AL-O-13 | GTCAGATATATACATCCTGTGTTCTACACCGAGTTCATCCCTTATGTGATGGACTC | r6-197 |
| AL-O-14 | AAGCGACGATAATAGTCATTGAGGTTG | forward-left-junction-YMR/601 |
| AL-O-15 | GTCTCCAGGCACGTGTCAGATATATACATCCATTAATTAATCCCCTTCAGCACGCAGC | reverse-left-junction-YMR/601 |
| AL-O-16 | GTCTACGACCAATTGAGCGGCCTCGGCACCGGGATTCTCCAGACAAGGTTTCTCACTATCAAGATGTACTGG | forward-right-junction-601/YMR |
| AL-O-17 | TACTCTTCAAGATCTAGAGGTTCGGGAAG | reverse-right-junction-601/YMR |
| AL-O-18 | CGGCCGCGTATCGATCATTTATCTTTCACTGC | p426-PCR-ClaI |
| AL-O-19 | CGGCCGCCGGTACCCAATTCGCCCTATAG | p426-PCR-KpnI |
| AL-O-20 | CGGCCGATCGATTGTGGGAAGTCGGCGCGACAGTTTTAGAGCTAGAAATAGCAAGT | gRNA-YMR262-cassette-forward |
| AL-O-21 | GGCGAATTGGGTACCGGCCGC | gRNA-cassette-reverse |
| AL-O-22 | GCCAGGACCTGCAATAACGTTTGTAC | foward-sequencing-left-junction-YMR/601 |
| AL-O-23 | GGGCATCTCCCTCTTCTGGGTCC | reverse-sequencing- |

| | | |
|---|---|---|
| | | right-junction-601/YMR |
| AL-O-24 | GAGCAGCTGCAGTATTTGAATGCAC | forward-southern-genomic-probe |
| AL-O-25 | GCTACCCGCACCGTATACAAAAAGG | Reverse-southern-genomic-probe |
